# Nuclear cholesterol is required for transcriptional repression by BASP1

**DOI:** 10.1101/2021.01.24.427800

**Authors:** Amy E. Loats, Samantha Carrera, Anna F. Fleming, Abigail R.E. Roberts, Alice Sherrard, Eneda Toska, Kathryn F. Medler, Stefan G. E. Roberts

## Abstract

Lipids are present within the cell nucleus where they engage with factors involved in gene regulation. Cholesterol associates with chromatin in vivo and stimulates nucleosome packing in-vitro, but its effects on specific transcriptional responses are not clear. Here we show that the lipidated WT1 transcriptional corepressor, BASP1, interacts with cholesterol in the cell nucleus through a conserved cholesterol interaction motif. We demonstrate that BASP1 directly recruits cholesterol to the promoter region of WT1 target genes. Mutation of BASP1 to ablate its interaction with cholesterol or the treatment of cells with drugs that block cholesterol biosynthesis inhibit the transcriptional repressor function of BASP1. We find that the BASP1-cholesterol interaction is required for BASP1-dependent chromatin remodelling and the direction of transcription programs that control cell differentiation. Our study uncovers a mechanism for gene-specific targeting of cholesterol where it is required to mediate transcriptional repression.

**Significance:** Cholesterol is present within the cell nucleus where it associates with chromatin but to date, a direct role for cholesterol in nuclear processes has not been identified. We demonstrate that the transcriptional repressor BASP1 directly interacts with cholesterol within the cell nucleus through a consensus cholesterol interaction motif. BASP1 recruits cholesterol to the promoter region of target genes where it is required to mediate chromatin remodelling and transcriptional repression. Our work demonstrates that nuclear cholesterol plays a direct role in transcriptional regulation.

## Introduction

It has been known for several decades that lipids and the enzymes that regulate lipid metabolism are present in the nucleus (Garcia-Gil et al., 2017; Fernandes et al., 2018; Fiume et al., 2019). The pool of nuclear lipids includes phospholipids, sphingolipids, gangliosides and cholesterol that associate with nuclear proteins, chromatin and the recently described nuclear lipid droplets (Barbosa and Siniossoglou, 2020). Several studies have reported that nuclear lipids associate with components of the transcription machinery but their function(s) in gene regulation are not clear (Garcia-Gil et al., 2017; Fernandes et al., 2018; Fiume et al., 2019).

BASP1 is a transcriptional cofactor that associates with WT1, converting it from a transcriptional activator to a repressor (Toska and Roberts, 2014). N-terminal myristoylation of BASP1 is required for its function as a transcriptional repressor and it uses this motif to interact with and recruit phosphatidylinositol 4,5-bisphosphate (PIP2) to gene promoters (Toska et al., 2012, 2014). The promoter-bound BASP1-PIP2 complex supports the assembly of chromatin remodelling proteins including Histone Deacetylase 1 (HDAC1) and BRG1 to WT1 target genes (Toska et al., 2012, 2014). BASP1 can also repress the function of other transcriptional activators, including c-myc (Hartl et al., 2009, 2020) and estrogen receptor α (Marsh et al., 2017), suggesting a broad role for this lipidated cofactor in the regulation of transcription.

WT1 and BASP1 cooperate to drive transcription programs that control differentiation in several model cell line systems (Belali et al., 2020; Essafi et al., 2012; Goodfellow et al., 2011; Green et al., 2008; Toska et al., 2012, 2014). Recent studies in mice found that knockout of BASP1 in taste receptor cells leads to loss of differentiated phenotype and the reactivation of WT1 target genes that are normally associated with the progenitor cells (Gao et al., 2019). A fundamental role for WT1-BASP1 in promoting the differentiated state was demonstrated with the finding that depletion of BASP1 in adult fibroblasts replaces the requirement of Sox2, c-Myc or Oct4 in the induction of fibroblasts to pluripotency (Blanchard et al., 2017). In these cells, BASP1 normally represses WT1 target genes and blocks pluripotency in favor of cell maintenance in their differentiated state.

BASP1 contains a functional cholesterol recognition consensus (CRAC) motif adjacent to its N-terminal myristoyl moiety and binds to cholesterol within cell membranes (Epand et al., 2004, 2005; Figure 1A). In this study we demonstrate that BASP1 interacts with nuclear cholesterol via its CRAC motif, recruiting it to the promoter region of WT1 target genes. We find that the BASP1-cholesterol interaction is required for transcriptional repression and that it plays a role in BASP1-dependent differentiation in-vitro. Our results demonstrate direct gene-specific control by nuclear cholesterol through interaction with the lipidated transcriptional repressor BASP1.

**Figure 1.**
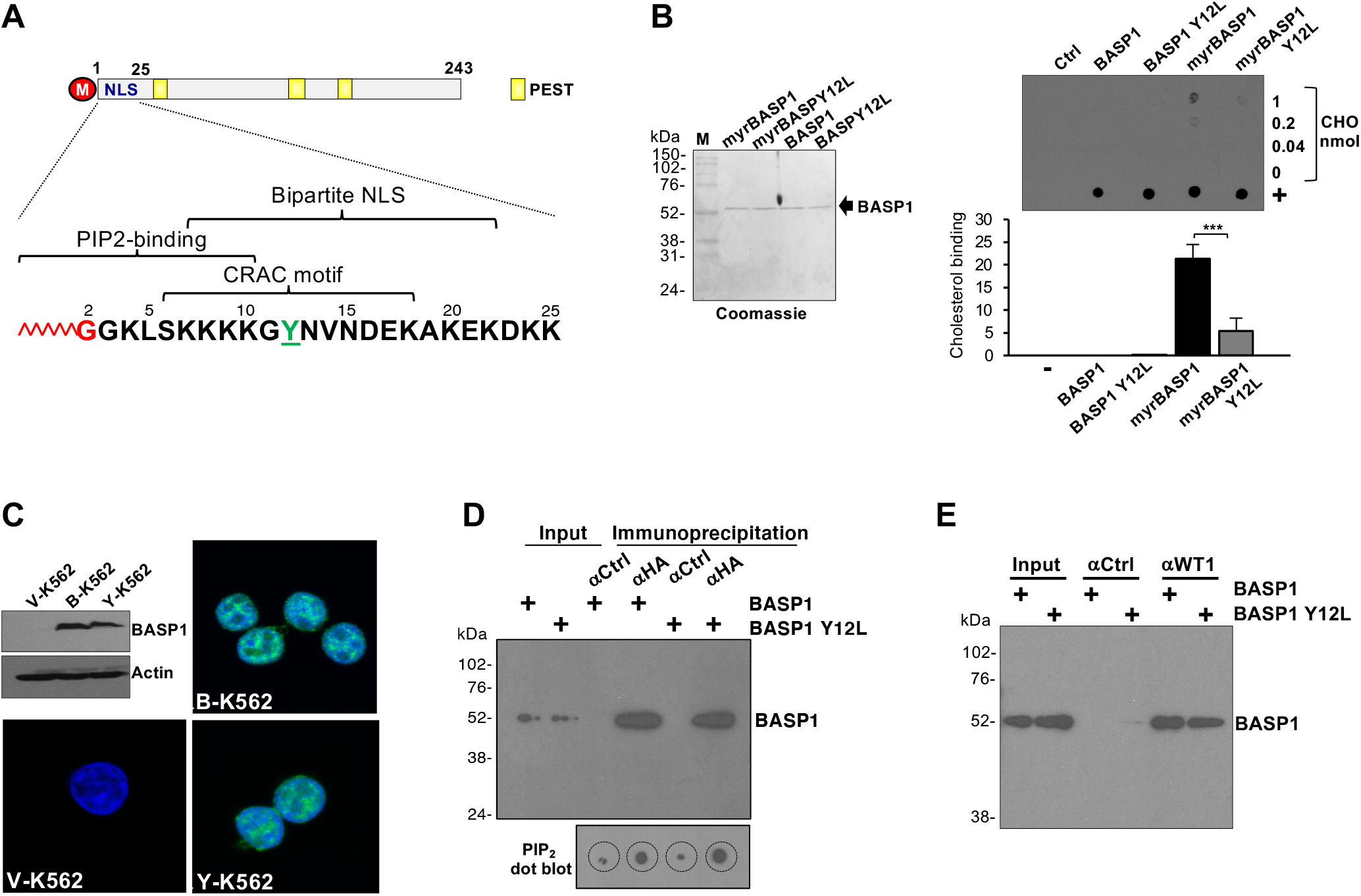
BASP1 interacts with cholesterol via a CRAC motif. (A) Schematic of BASP1 with N-terminal myristoyl (red), nuclear localization sequence (NLS) and PEST motifs. The N-terminal 25 residues, myristoylated G2 (red), PIP2-binding domain, bipartite NLS and CRAC motif (central Tyr underlined green) is below. (B) Myristoylated and non-myristoylated wtBASP1 or BASP1-Y12L prepared from *E.coli* (coomassie gel at left) were used to probe membranes dot blotted with cholesterol (0.04, 0.2 and 1nmol). + is a positive control for BASP1 binding (sheep BASP1 antibody). Ctrl is a membrane probed with BSA while the other panels were probed with the BASP1 derivatives and detected with rabbit BASP1 antibody. Image J quantitation is below for three independent experiments. Error bars are standard deviation of the mean (SDM) of three independent experiments. *** is students t-test p<0.001. (C) Immunoblot of whole cell extracts prepared from V-K562, B-K562 or Y-K562 cells. Immunofluorescence shows that wtBASP1 and BASP1-Y12L are nuclear in K562 cells. (D) Control (αCtrl) or HA (αHA) antibodies were used for BASP1 immunoprecipitation with nuclear extracts from B-K562 or Y-K562 cells and probed for BASP1 by immunoblot and for PIP2 by dot blot (below). Input is 2% of the nuclear extract used in each immunoprecipitation. (E) Control IgG (αCtrl) or WT1 (αWT1) antibodies were used for immunoprecipitation with nuclear extracts derived from B-K562 or Y-K562 cells and the content probed for BASP1 by immunoblot. Input is 5% of the nuclear extract used in each immunoprecipitation.

## Results

### BASP1 interacts with cholesterol in the cell nucleus through a conserved CRAC motif

BASP1 interacts with cholesterol-rich domains (Epand et al., 2004, 2005) and contains a consensus CRAC motif (L/V-X(1-5)-Y-X(1-5)-R/K) located within the N-terminus close to the myristoyl motif (Fantini et al., 2019; Figure 1A). Using *E.coli* engineered to express N-myristoyl transferase (NMT), we generated recombinant myristoylated wild type (wt) BASP1 containing a C-terminal His-tag (myrBASP1) and the myristoylated mutant derivative Y12L (myrBASP1 Y12) that ablates the critical tyrosine residue of the CRAC motif. We also expressed the same proteins in *E.coli* lacking NMT to produce non-myristoylated wtBASP1 and BASP1 Y12L. The purified proteins are shown in Figure 1B. In filter binding assays, recombinant myristoylated wtBASP1 bound directly to cholesterol but the non-myristoylated forms of BASP1 did not. Moreover, the interaction of myristoylated BASP1-Y12L with cholesterol was significantly reduced (Figure 1B, see quantitation below). Thus, both myristoylation of BASP1 and an intact CRAC motif are required for the direct interaction with cholesterol.

Our previous work has used K562 chronic myelogenous leukaemia (CML) cells which lack endogenous BASP1 but contain endogenous WT1. Stably introducing wtBASP1 into K562 cells (B-K562) represses WT1-dependent target genes (Goodfellow et al., 2012; Toska et al., 2012, 2014). We generated a K562 cell line derivative that expresses BASP1-Y12L (Y-K562) which, like wtBASP1, is largely nuclear (Figure 1C; V-K562 are vector control cells). Immunoprecipitation of BASP1 from nuclear extracts via a C-terminal HA-tag followed by dot-blotting to detect PIP2 confirmed that wtBASP1 binds PIP2 (Figure 1D; Toska et al., 2012, 2014). BASP1 Y12L immunoprecipitates contained a similar level of PIP2 indicating that the BASP1-PIP2 interaction does not depend on the interaction of BASP1 with cholesterol. BASP1 Y12L also co-immunoprecipitated with WT1 from nuclear extracts at a level comparable with wtBASP1 (Figure 1E).

We next sought to determine if BASP1 interacts with cholesterol in the cell nucleus. To resolve nuclear cholesterol from cytoplasmic cholesterol, we isolated nuclei with low cytoplasmic contamination as determined by levels of the golgi protein TGN46 (Figure S1A) and the ER marker Calreticulin (Figure S1B). The K562 cell line derivatives were incubated in lipid-free media supplemented with a 27-alkyne cholesterol derivative that is utilized by cells as standard cholesterol (Hofmann et al., 2014). Nuclei were then prepared and using click chemistry, the 27-alkyne cholesterol was crosslinked to azide-PEG3-biotin which was then detected using a streptavidin-linked fluorophore. BASP1 was simultaneously probed using immunocytochemistry. Cholesterol was detected within the purified nuclei of K562 cells (Figure 2A; CHO) and showed reduced colocalisation with BASP1-Y12L compared to wtBASP1 (Manders quantitation of the proportion of BASP1 signal that overlaps with cholesterol is shown below, left). Endogenous BASP1 in MCF7 cells also colocalized with 27-alkyne cholesterol within the nucleus (Figure S1C). We next used click chemistry combined with immunoprecipitation to test the interaction between BASP1 and nuclear cholesterol. V-K562, B-K562 and Y-K562 cells were cultured with 27-alkyne cholesterol as above for 48 hours (as above) and then underwent the click chemistry reaction with azide-PEG3-biotin. Nuclear protein extracts were prepared followed by precipitation with streptavidin-coated beads. wtBASP1, but not BASP1-Y12L, co-precipitated with alkyne-cholesterol (Figure 2B). Similar analysis verified that 27-alkyne cholesterol associates with endogenous BASP1 in the nuclei of MCF7 cells (Figure 2C). Taken together, the data in Figures 1 and 2 demonstrate that BASP1 interacts with nuclear cholesterol via a functional CRAC motif.

**Figure 2.**
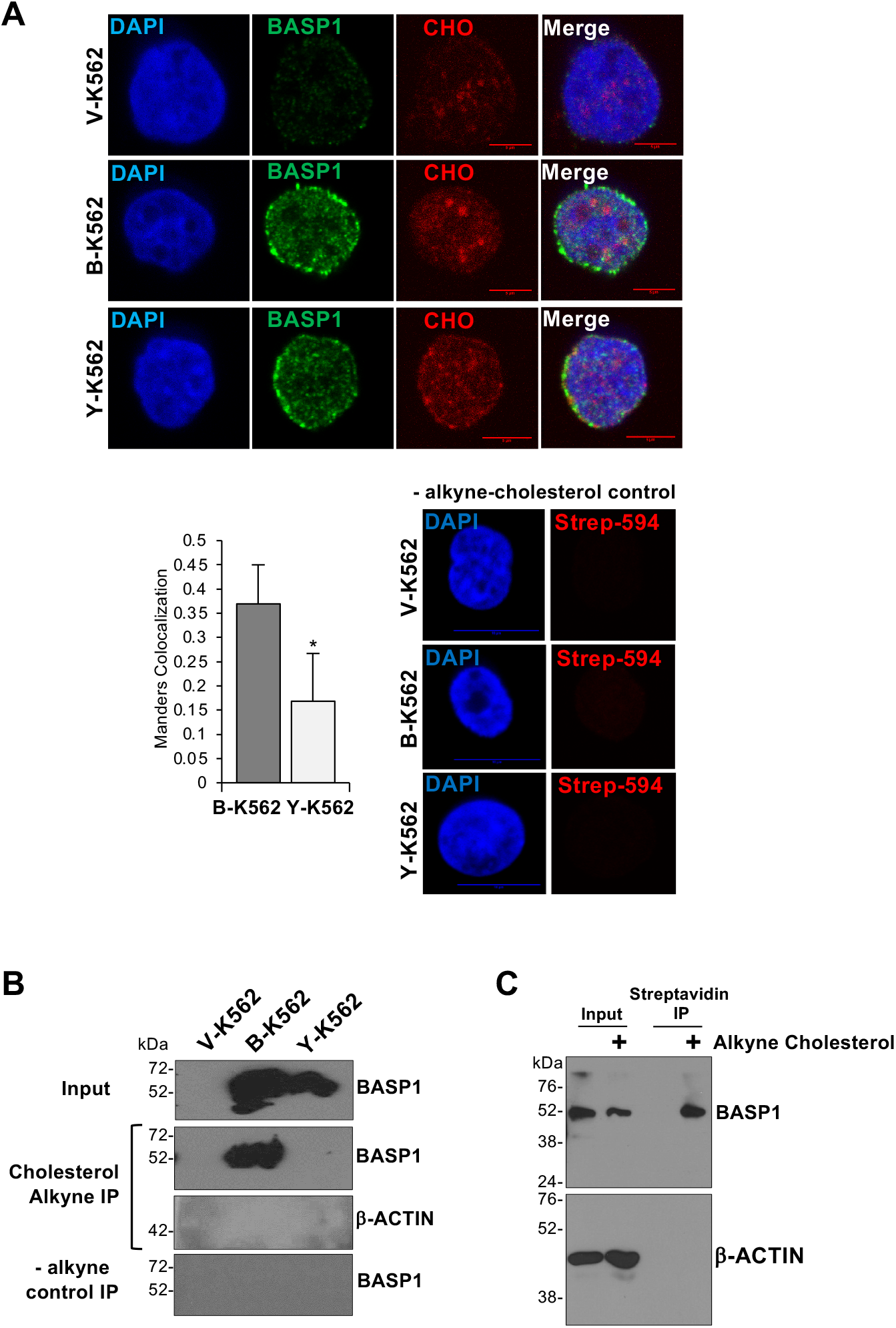
BASP1 interacts with nuclear cholesterol. (A) K562 cell derivatives were incubated with 10mg/ml alkyne cholesterol (CHO) in lipid-free media for 20 hours. Nuclei were purified and after treatment with azide PEG3-biotin and click reaction using 2mM CuBF4, subjected to immunocytochemistry with streptavidin-linked fluorophore (CHO) or BASP1 antibodies. Scale bar is 5μm. Below right, cells treated in the same way, but without CHO-alkyne. Manders analysis to quantitate the colocalisation of cholesterol with wtBASP1 and BASP1-Y12L (below, left). Error bars are SDM of three independent experiments. * is students t-test p<0.05. (B) As in part A but nuclear extract was prepared and precipitation performed with streptavidin magnetic beads then immunoblotted with BASP1 or actin antibodies. -alykne control IP is the same click chemistry reaction but with cells not incubated with alkyne cholesterol. (C) MCF7 cells were incubated with (+) or without (−) alkyne cholesterol, nuclear extract prepared followed by click chemistry and the alkyne cholesterol captured with streptavidin magnetic beads. The content was probed with BASP1 or β-actin antibodies. Input is 5% of the nuclear extract used in each assay.

### BASP1 recruits cholesterol to the gene promoter region of WT1 target genes

We next determined if an intact CRAC motif in BASP1 is required for transcriptional repression. As we have previously shown, wtBASP1 repressed transcription of the WT1 target genes AREG, ETS-1, JUNB and VDR (Goodfellow et al, 2011; Toska et al., 2012, 2014; Figure 3A). In contrast, BASP1 Y12L was defective in transcriptional repression. ChIP analysis determined that both wtBASP1 and BASP1 Y12L were recruited to the promoter region of all the genes tested (Figure 3B). Combining click chemistry with ChIP (Click-ChIP) we found that cholesterol associates with the AREG, ETS-1, JUNB and VDR promoters at a significantly increased level when wtBASP1 is also present at the promoter (Figure 3C; compare V-K562 with B-K562). In K562 cells that express BASP1-Y12L, cholesterol was recruited at a significantly reduced level at the AREG, JUNB and VDR promoters compared to B-K562 cells and, although not statistically significant, was also reduced at the ETS-1 promoter. Thus, BASP1 recruits cholesterol to the promoter region of WT1 target genes and this requires an intact CRAC motif in BASP1.

**Figure 3.**
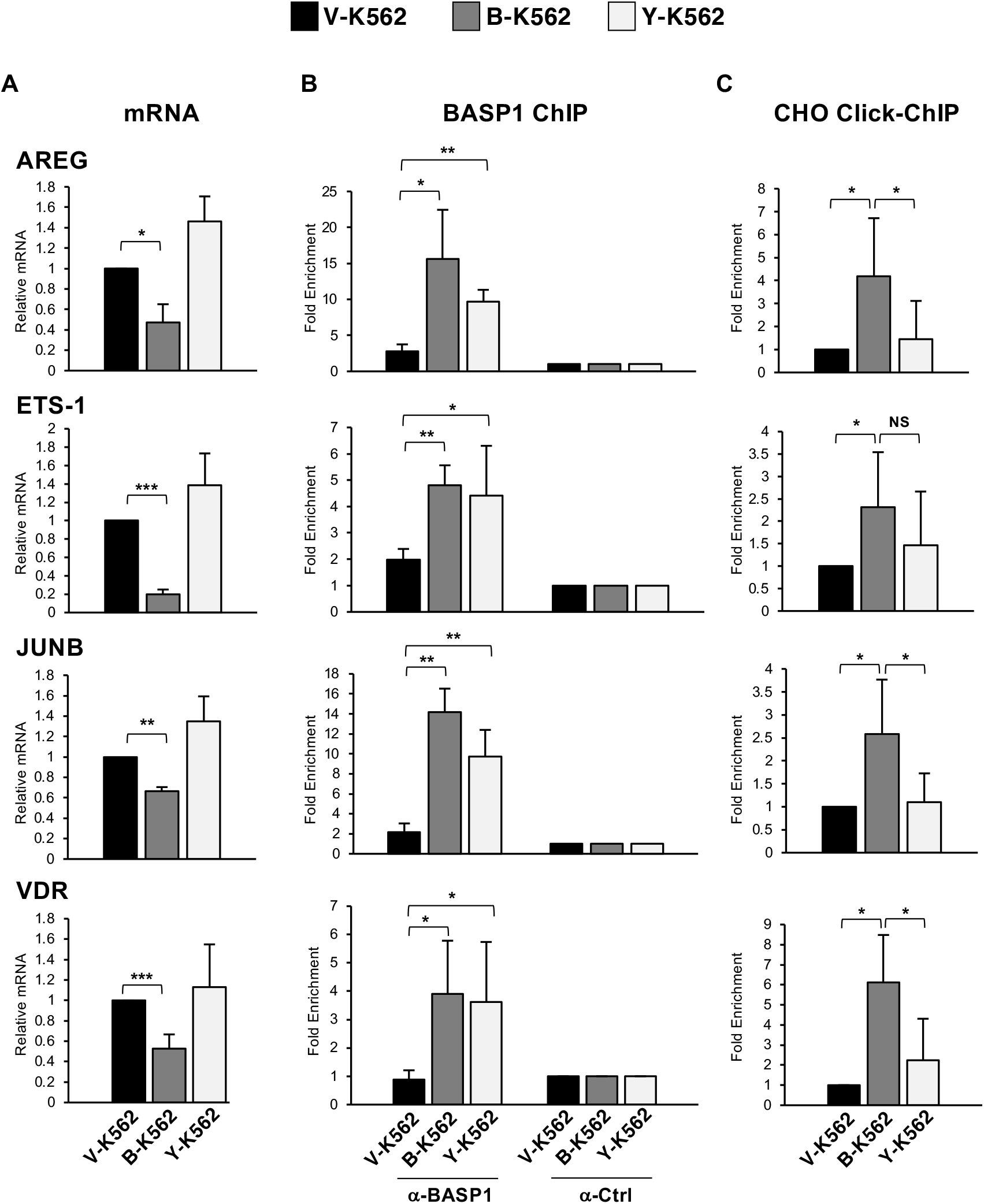
BASP1 Y12L is defective in transcriptional repression and fails to recruit cholesterol to the promoter of WT1 target genes. (A) cDNA was prepared from V-K562, B-K562 or Y-K562 cells. Expression of AREG, ETS-1, JUNB and VDR were compared to control GAPDH. (B) Cells as above were subjected to ChIP analysis with αBASP1 or control (αCtrl) IgG. Presented as fold enrichment over a control genomic region. (C) K562 cell lines were incubated with alkyne cholesterol as in Figure 1E. After the click chemistry reaction the cells were subjected to ChIP with streptavidin linked beads (Click-ChIP). Results are fold-enrichment over a control genomic region. Error bars are SDM from at least three independent experiments. Students t-test; *<0.05, **<0.01, ***<0.001, not significant (NS).

### The BASP1-cholesterol interaction is required for the control of differentiation

A central function of the WT1-BASP1 complex is to drive cell differentiation programs (Toska and Roberts, 2014). BASP1 cooperates with WT1 to divert the phorbol ester-induced differentiation program of K562 cells away from megakaryocytes to neuronal-like cells in both morphology and function (Goodfellow et al., 2019; Toska et al., 2012, 2014). This was driven by the WT1/BASP1-dependent downregulation of genes that specify megakaryocyte identity accompanied by the upregulation of neuronal markers. We tested if BASP1 Y12L was able to support the neuronal-like differentiation program of K562 cells. The K562 cell derivatives were induced to differentiate with PMA for 72 hours and the extent of cellular arborization was visualized by phase microscopy (Figure 4A, quantitation shown below). wtBASP1 expression in K562 cells led to induction of an arborized phenotype compared to V-K562 cells that was significantly reduced in Y-K562 cells.

**Figure 4.**
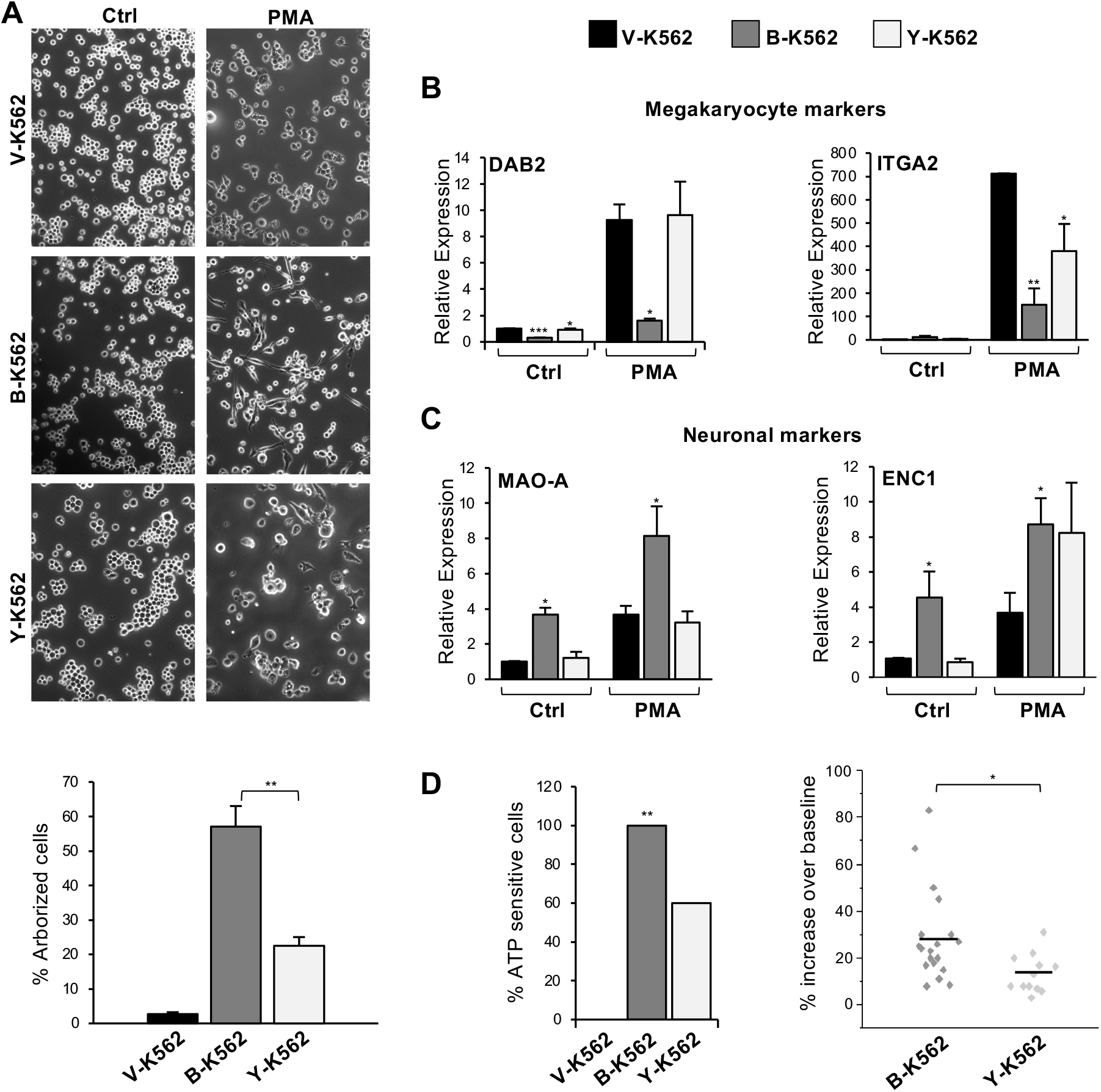
An intact CRAC motif in BASP1 is required to induce a neuronal-like differentiation program in K562 cells. (A) V-K562, B-K562 or Y-K562 cells were treated with 200nM PMA or DMSO for 72 hours and then observed by phase microscopy. Percentage arborized cells is below. Error bars are SDM of three independent experiments and ** indicates students t-test p<0.01. (B) Cells were treated as in part A but 48 hours later cDNA prepared and qPCR used to determine expression of the megakaryocyte markers DAB2 and ITGA2 relative to GAPDH. Error bars are SDM of three independent experiments with students t-test; *p<0.05, **p<0.01, ***p<0.001. (C) As in part B except that neuronal markers MAOA and ENC1 were analyzed. (D). The K562 derivatives were treated as in part A, ATP (10 mM) was applied and calcium influx measured by fluorescence imaging. At left, percentage of cells that elicited a calcium influx response. * indicates chi squared test at p<0.01. At right, the responsive B-K562 cells (n=19 cells) and Y-K562 cells (n=12 cells) are shown as a scatter plot against the amplitude of response. * indicates p<0.05 by students t-test.

Analysis of the level of mRNA encoding the megakaryocyte-specific markers DAB2 and ITGA2 revealed that whereas wild type BASP1 blocked their induction as we reported before (Goodfellow et al, 2011), BASP1 Y12L failed to repress transcription of DAB2 and was defective in repression of ITGA2 (Figure 4B). Analysis of neuronal-specific markers (MAO-A and ENC1) showed that BASP1 induced their expression as we previously reported (Figure 4C; Goodfellow et al., 2011). BASP1 Y12L was defective in inducing MAO-A but still able to support induction of ENC1. These data suggest that BASP1 Y12L is selective in its effects in diverting the PMA-induced K562 cell differentiation program to neuronal-like cells. We next determined the response of the PMA-treated cells to the neurotransmitter ATP. As we reported previously (Goodfellow et al., 2011), none of the control V-K562 cells mobilized Ca^2+^ stores in response to addition of ATP, but 100% of the B-K562 cells did (Figure 4D, left). Y-K562 cells showed an intermediate response (60% of the cells). Analysis of the size of the Ca^2+^ signals in the B-K562 and Y-K562 cells that responded found reduced Ca^2+^ mobilization in BASP1 Y12L cells compared to wtBASP1 cells (Figure 4D, right). Thus, the CRAC motif in BASP1 plays a role in the diversion of the differentiation program of K562 cells to a neuronal-like cell.

### Inhibition of cholesterol synthesis abolishes transcriptional repression by BASP1

Our results so far have shown that mutation of the critical tyrosine residue of the BASP1 CRAC motif disrupts the interaction with cholesterol, its recruitment to the promoter and causes a defect in transcriptional repression and BASP1-induced differentiation. To confirm and extend these findings we determined the effect of cholesterol-depleting drugs on BASP1 function. We first confirmed that the drugs were used at an effective concentration to significantly reduce nuclear cholesterol (Figure S2A). We treated V-K562 cells or B-K562 cells with Atorvastatin, an inhibitor of HMG-CoA reductase (the rate-limiting enzyme in cholesterol biosynthesis) and determined the effect on BASP1-mediated transcriptional repression. Atorvastatin treatment abolished transcriptional repression of AREG and ETS-1 by BASP1 in K562 cells (Figure 5A). Similar experiments with Lovastatin (which also inhibits HMG-CoA reductase) produced comparable results (Figure S2B). We also tested the effect of Atorvastatin on BASP1 transcriptional repressor function in MCF7 cells transiently transfected with either a control siRNA or siRNA targeting BASP1. Knockdown of endogenous BASP1 in MCF7 cells led to relief of repression of WT1 target genes as previously reported (Figure 5B; Toska et al., 2014). When the cells were treated with atorvastatin, the relief of transcriptional repression upon BASP1 knockdown did not occur. We observed similar effects when MCF7 cell line derivatives that stably express either a control shRNA (shNEG) or BASP1 shRNA (shBASP1, targeting a different sequence to the siRNA used in Figure 5B) were treated with Lovastatin (Figure S2C). Statins inhibit the biosynthesis of both cholesterol and isoprenoids (Mullen et al, 2016). We therefore next tested the effect Triparanol, an inhibitor of 24-dehydrocholesterol reductase, which catalyzes the final step in cholesterol biosynthesis pathway and does not inhibit isoprenoid production. As we had observed with the statins, Triparanol blocked the transcriptional repressor function of BASP1 (Figure 5C). We next determined if interference with cholesterol biosynthesis in K562 cells affects the BASP1-dependent differentiation to neuronal-like cells. As before, treatment of B-K562 cells, but not V-K562 cells, with PMA induced an arborized phenotype (Figure 5D). This was blocked if the cells were simultaneously treated with Atorvastatin (see quantitation below). Taken together, the data in Figures 5A-D demonstrate that inhibition of cholesterol biosynthesis blocks the transcriptional repressor function of BASP1 and impedes BASP1-induced differentiation.

**Figure 5.**
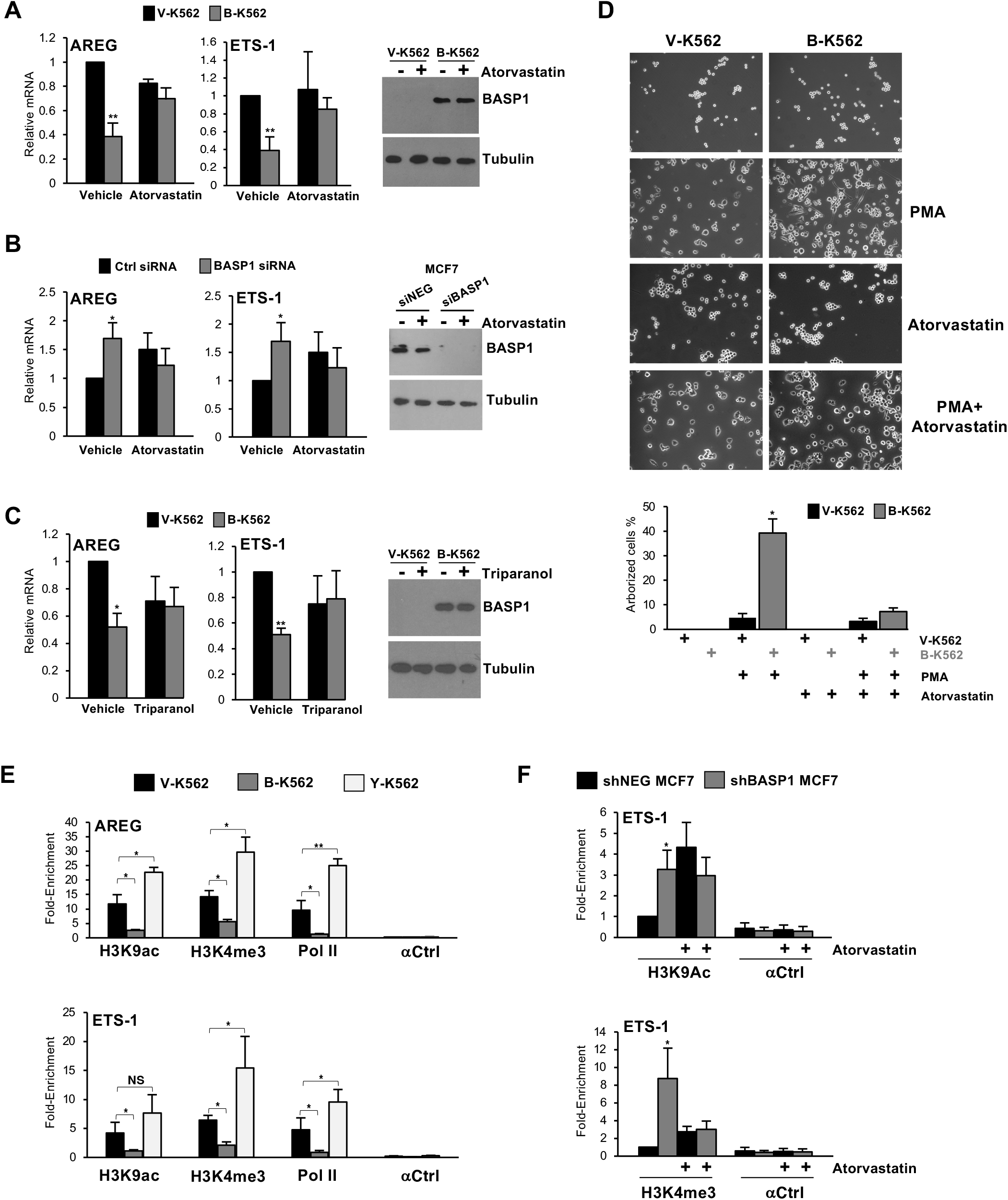
Statins specifically inhibit the transcriptional repressor function of BASP1. (A) V-K562 or B-K562 cells were treated with 20μM Atorvastatin or drug vehicle (DMSO) for 48 hours. cDNA was prepared and expression of AREG and ETS-1 was determined relative to GAPDH. At right, a western blot of BASP1 is shown. (B) MCF7 cells were transfected with either a control siRNA (siNEG) or siRNA targeting BASP1 (siBASP1). 24 hours later the cells were treated with 20μM Atorvastatin or drug vehicle (DMSO) for 48 hours. cDNA was prepared and expression of AREG and ETS-1 was determined relative to GAPDH. At right, a western blot of BASP1 is shown. (C) As in part A except that cells were treated with 20μM Triparanol. (D) V-K562 cells or B-K562 cells were treated with PMA (200nM) and/or Atorvastatin (20μM) as indicated. 72 hours later the cells were analyzed as in Figure 3A. (E) ChIP was performed with V-K562, B-K562 and Y-K562 cells with control IgG (αCtrl), H3K9Ac, H3K4me3 and RNA polymerase II (Pol II) antibodies. Data are presented as fold-enrichment at the AREG (top) and ETS-1 (bottom) promoters over a control genomic region. (F) MCF7 cells stably expressing either control shRNA (shNEG) or BASP1 shRNA (shBASP1) were treated with 20μM Atorvastatin or vehicle (DMSO). 48 hours later the cells were crosslinked and ChIP performed with antibodies against H3K9ac (top graph) or H3K4me3 (bottom graph) alongside IgG control (αCtrl). Data are presented as fold enrichment at the ETS-1 promoter against a control genomic region. Error bars are SDM of at least three independent experiments (*p<0.05 and **p<0.01 and NS is not significant by students t-test).

### The BASP1-cholesterol interaction is required for chromatin remodelling

BASP1 acts as a transcriptional repressor by remodeling chromatin and blocking the recruitment of RNA polymerase II at the promoter region of WT1 target genes (Pol II; Toska et al., 2012, 2014; Essafi et al., 2014). We next performed ChIP analysis with the K562 cell line derivatives to determine the effect of wtBASP1 and BASP1-Y12L on Pol II recruitment as well as two histone modifications indicative of transcriptional activity (H3K9Ac and H3K4me3) at the AREG and ETS-1 promoters (Figure 5E). Compared to V-K562 cells, B-K562 cells show significantly reduced H3K9Ac, reduced H3K4me3, and reduced Pol II binding at the AREG and ETS-1 promoters. BASP1 Y12L did not support the reduction in either the active chromatin marks or in Pol II recruitment. Indeed, there was a significant increase over the vector control cells suggesting the possibility that BASP1 Y12L has transcriptional activator properties. We next performed ChIP analysis using MCF7 cell line derivatives that express either a control shRNA (sh-NEG MCF7) or an shRNA that targets BASP1 (shBASP1) to determine the effect of Atorvastatin on H3K9Ac and H3K4me3 marks at the ETS-1 promoter. Consistent with the observations in K562 cells, shRNA-mediated knock down of BASP1 in MCF7 cells caused an increase in H3K9Ac and H3K4me3 marks at the ETS-1 promoter (Figure 5F). However, treatment of the MCF7 cell line derivatives with Atorvastatin abolished the BASP1-dependent changes in these marks. Taken together, the data in Figures 5E and F suggest that the BASP1-cholesterol interaction is required for BASP1-dependent histone modification.

## Discussion

In this study we provide evidence for the directed recruitment of nuclear cholesterol to gene promoters by BASP1 and that the BASP1-cholesterol interaction is required for BASP1-dependent transcriptional repression. Previous work has reported that cholesterol binds directly to histones and drives the compaction of nucleosomes in-vitro (Silva et al., 2017; Fernandes et al., 2018). Cholesterol interacts with at least six sites on the nucleosome and is proposed to facilitate dewetting of hydrophobic surfaces on the histones, leading to enhanced histone-histone contacts that drive chromatin condensation. Thus, BASP1-mediated recruitment of cholesterol might elicit direct effects on the nucleosome that contribute to establishing transcriptionally repressive condensed chromatin at specific genes. Cholesterol binding to BASP1 may also facilitate the recruitment of chromatin remodelling activities. Previous studies found that the BASP1-PIP2 interaction is required to mediate contact between BASP1 and HDAC1 (Toska et al., 2012). Thus, the BASP1-cholesterol interaction might similarly facilitate the recruitment of chromatin remodelling complexes. In support of this notion, we find that a mutation of the BASP1 CRAC motif ablates the BASP1 dependent deacetylation of H3K9 and demethylation of H3K4 at target gene promoters.

A role for lipids in regulating the affinity and/or specificity of protein-protein interactions in transcription regulation is reminiscent of the action of lipid-derived hormones on steroid receptor transcription factors (Sever and Glass, 2013). However, the widely reported roles of nuclear lipids in transcription suggests a more general function in transcription control. The regulation of transcription by lipids might also have additional physiological importance. Knockout of WT1 in adult mice leads to rapid fat loss through direct effects on adipocyte homeostasis and lipid metabolism (Chau and Hastie, 2015). Indeed, several genes in the cholesterol biosynthesis pathway and lipid transport are target genes of WT1 (Rae et al., 2004; Tamura et al., 2020). It is possible that the control of WT1-BASP1 by lipids provides a feedback mechanism to regulate lipid metabolism.

How lipids are sequestered within the nucleus and the regulation of their subnuclear localization is not yet clear but lipidated transcription factors such as BASP1 can provide at least part of the mechanism. Nuclear lipid droplets (NLDs) have recently been described that may play a role in the regulation of transcription factor accessibility (Welte, 2015; Farese and Walther, 2016; Barbosa and Siniossoglou, 2020). BASP1 is a component of cytoplasmic cholesterol-rich lipid droplets (Khor et al., 2014), which raises the prospect that BASP1 might also be incorporated into cholesterol-rich NLDs. NLDs associate with PML bodies (Ohsaki et al., 2016) which can also contain BASP1 (Green et al., 2008). It is therefore possible that BASP1-lipid interactions may control transcription processes through the formation of phase separated nuclear bodies (Sztacho et al., 2019). Lipidated transcription factors like BASP1 have the potential to play important roles in liquid-liquid phase separation in chromatin remodelling and transcription control (Erdel and Rippe, 2018; Wang et al., 2019).

## Materials and Methods

### DNA constructs

pcDNA3 BASP1-HA is described in Goodfellow et al. (2011). pcDNA3-BASP1 Y12L-HA was generated using the Quickchange site-directed mutagenesis kit (Agilent) and verified by DNA sequencing. The coding sequence of wtBASP1 and BASP1 Y12L lacking the stop codon was amplified and cloned in-frame into the NcoI/XbaI sites of pET23d bacterial expression vector to generate BASP1 and BASP1 Y12L with a C-terminal His-tag. pRSF-1-NMT for expression of N-myristoyl transferase in bacteria was purchased from Addgene.

### Cell lines, transfection and antibodies

K562 cells were maintained in RPMI 1640 (ThermoFisher) supplemented with 10% (v/v) fetal calf serum (Sigma-Aldrich), 1% (v/v) Pen-Strep (Sigma-Aldrich) and 1% (v/v) L-glutamine (Sigma-Aldrich). MCF7 cells were maintained in DMEM (ThermoFisher) supplemented with 10% (v/v) fetal calf serum and 1% (v/v) Pen-Strep. Stably transfected K562 and MCF7 cell lines were additionally supplemented with 1mg/ml G-418 (Sigma-Aldrich). All cells were kept at 37°C in humidified 95% air and 5% CO_2_ atmosphere. Cells were transfected with plasmids using Effectene (Qiagen) and with siRNA using HiPerFect (Qiagen). Antibodies used are listed in Supplementary Table 1.

### Chromatin Immunoprecipitation

K562 cells were collected and resuspended to a density of 1×10^6^ cells/ml in PBS. Crosslinking was achieved by adding formaldehyde to a final concentration of 1.42% (v/v), and incubating at room temperature for 15 mins. Formaldehyde was quenched with ice-cold 125mM glycine for 5 mins at room temperature. Cells were collected by centrifugation at 2000 × g for 5 mins at 4°C and washed with ice cold PBS. Cells were resuspended to a concentration of 1×10^7^ cells/ml, and lysed in 1ml IP buffer (150mM NaCl, 50mM Tris-HCl (pH 7.5), 5mM EDTA, 0.5% (v/v) NP-40 and 1% (v/v) Triton X-100) plus protease inhibitor cocktail for 15 mins on ice. Lysed samples were centrifuged at 2000 × g for 5 mins at 4°C, and the pellet resuspended in 1ml IP buffer plus protease inhibitors for sonication. Chromatin was sheared via sonication using a QSonica Q500 at 60% amplitude. Successful sonication giving fragments 200-500bps in length was confirmed by resolving a small (decrosslinked) sample on a 1.5% (w/v) agarose gel. Following sonication the lysate was cleared by centrifugation at 12000 × g for 10 mins at 4°C. Samples were precleared by incubating with 10μl Protein G magnetic beads (ThermoFisher) for 1 hour, rotating at 4°C. 5μl Protein G magnetic beads, 600μl IP buffer, 1μl 10mg/ml acetylated BSA and appropriate antibody were also incubated together in one microtube per desired chromatin immunoprecipitation for a minimum of 4 hours, rotating at 4°C. 200μl of pre-cleared chromatin was then added to the incubated microtube of antibody and beads and rotated at 4°C overnight. A 2% input sample was also stored for later decrosslinking and processing.

Immunoprecipitated samples were then magnetised, supernatant discarded and beads sequentially washed once in IP buffer, high salt IP buffer (500mM NaCl, 50mM Tris-HCl (pH 8.0), 5mM EDTA, 0.5% (v/v) NP-40, 1% (v/v) Triton × 100), LiCl buffer (10mM Tris-HCl (pH 8.0), 250mM LiCl, 1mM EDTA, 1% (v/v) NP-40, 1% (w/v) Sodium Deoxycholate), and TE buffer (10mM Tris-HCl (pH 8.0), 1mM EDTA). After the final wash beads and 2% input samples were resuspended in 100μl of PK buffer (125mM Tris-HCl (pH 8.0), 10mM EDTA, 150mM NaCl, 1% (w/v) SDS) and placed at 65°C overnight. Samples were then incubated with 1μl of 20mg/ml Proteinase K for 3-4 hours at 55°C. Finally, the immunoprecipitated DNA was purified using the Qiaquick PCR purification kit (Qiagen) according to manufacturer’s instructions. Eluted DNA was placed at 95°C for 10 mins and prepared for qPCR using the primers in Supplementary Table 2.

### Immunoprecipitation

All protein immunoprecipitation was carried out using nuclear extracts prepared from K562 cells. Cells were collected by centrifugation for 3 mins at 1400 × g. The packed cell volume (PCV) was estimated, and the pellet incubated with 2/3 PCV of NE1 buffer (10mM HEPES pH 8.0, 1.5mM MgCl_2_, 10mM KCL, 1mM DTT) for 15 mins on ice, with protease inhibitors. Cells were then passed through a 23-gauge needle 4-5 times, and microfuged at full speed for 30 secs to isolate nuclei. The supernatant was discarded and remaining pellet resuspended in one PCV of NE2 buffer (20mM HEPES pH 8.0, 1.5mM MgCl_2_, 25% (v/v) Glycerol, 420mM NaCl, 0.2mM EDTA, 1mM DTT and 0.5mM PMSF) on ice for with regular stirring for 30 mins, with protease inhibitor cocktail. Nuclear debris was pelleted by 5 mins microfuge at full speed. The supernatant (nuclear extract) was dialysed against Buffer D (20mM HEPES pH 8.0, 100mM KCl, 0.2mM EDTA, 20% (v/v) Glycerol and 0.5mM DTT) for 2 hours at 4°C. Dialysed samples were cleared by centrifugation at 13,000 × g for 5 mins at 4°C and precleared by incubation with 10μl Protein-G magnetic beads and appropriate IgG antibody for 1-2 hours at 4°C. Following the preclear, samples were divided as necessary onto fresh Protein G magnetic beads, IP buffer (20mM HEPES pH 8.0, 100mM KCl, 0.2mM EDTA, 20% (v/v) Glycerol, 0.5mM DTT and 0.05% (v/v) NP-40) and the appropriate antibody. Samples were rotated at 4°C overnight. The magnetic beads were collected and washed 3 times in 1ml IP buffer. For western blot, beads were resuspended in 10μl 1 × SDS loading buffer per gel and incubated at 95°C for 3 mins before SDS-PAGE and western blotting.

### Quantitative RT-PCR analysis

Total RNA was prepared using the RNeasy kit (Qiagen) and cDNA prepared using the Iscript cDNA synthesis kit (Bio-Rad). qPCR was performed using iQ SYBR green master mix (Bio-Rad) with samples run in triplicate on a Bio-Rad Miniopticon system. Melt curve analysis was performed at the end of each run. Data were collected using the BioRad-CFX Manager software. The relative abundance of each transcript was compared with control GAPDH. Primers used to amplify cDNA are listed in Supplementary Table 2.

### Immunofluorescence

K562 cells grown in suspension culture were collected via centrifugation at 1400 × g for 3 mins. Nuclei were isolated using the Nuclei EZ Prep nuclei isolation kit (Sigma-Aldrich). Nuclei were fixed in 4% (v/v) paraformaldehyde and rotated at room temperature for 15 mins, then incubated with 50mM NH_4_Cl for 15 mins, rotating, then washed three times in PBS. The nuclei were then incubated in blocking buffer (2% (w/v) BSA, 0.25% (w/v) Gelatin, 0.2% (w/v) Glycine, 0.2% (v/v) Triton X-100 in PBS) for 1 hour, rotating at room temperature. Primary antibody was diluted in PBS with 1% (w/v) BSA, 0.25% (w/v) gelatin and 0.2% (v/v) Triton X-100 and incubated for 1 hour, rotating at room temperature. Nuclei were washed three times with washing buffer (0.2% (w/v) Gelatin in PBS). Fluorescent secondary antibody was then applied for 45 mins in the same buffer as the primary antibody, and samples were rotated in the dark. Nuclei were washed three times in washing buffer before 10 mins incubation with DAPI solution. Nuclei were resuspended in a minimum volume of DABCO mounting media (Sigma-Aldrich) to mount onto poly-lysine coated slides. Samples were viewed using a Leica SP5-II AOBS confocal laser scanning microscope attached to a Leica DM I6000 inverted epifluorescence microscope with oil 63x lens. Images were processed using ImageJ or Volocity 6.3 software.

### Click Chemistry

K-562 and MCF7 cells were incubated in lipid free media (RPMI and DMEM, respectively) containing 10μg/ml alkyne-Cholesterol (Avanti) for 16 hrs. Samples were washed once in PBS and nuclei obtained using the EZ Prep nuclei kit then incubated with 50μM Azide-PEG3-biotin conjugate (Sigma-Aldrich) was dissolved in prewarmed Buffer A and added to samples.. The click reaction was initiated via addition of 2mM CuBF4 in 2% (v/v) acetonitrile, and the reaction was left to proceed at 43°C for 30 mins with gentle agitation. The nuclei were then washed in Buffer A and immediately used for either preparation of nuclear extract for immunoprecipitation (using streptavidin magnetic beads, ThermoFisher), chromatin immunoprecipitation (Click-ChIP with streptavidin magnetic beads) or immunofluorescence (using Streptavidin Alexa Fluor 594 conjugate, ThermoFisher).

### Protein production, cholesterol assay and cholesterol-binding assay

For production of BASP1 proteins, BL21 DE3 cells were initially transformed with pRSF-1-NMT for the expression of N-myristoyl transferase, then with the pET23d BASP1 expression vectors. 20ml of culture in LB medium was seeded overnight at 37°C, then added to 200ml of media and incubated for 1hour in a shaker at 37°C.

Where required, myristic acid was added to a final concentration of 0.2mg/ml. The optical density (at 600nm) was monitored and when OD600=0.6 IPTG (1mM) was added. After incubation a 37°C for 3 hours the cells were harvested and the pellet frozen at −80°C. The His-tagged BASP1 derivatives were prepared using Ni-NTA beads following the manufacturers instructions (Qiagen). Purified proteins were dialysed into Buffer D and stored at −80°C.

Cholesterol was solubilised in methanol/chloroform/water (2:1:0.8 v/v/v) at a final concentration of 1mM, diluted as required, and dotted onto PVDF membrane. The membrane was incubated in binding buffer (20mM Tris pH8, 300mM NaCl, 0.5% (v/v) Tween, 2.5% (w/v) dried milk). For 1 hour rocking at room temperature. Fresh binding buffer was added containing the BASP1 derivative at a concentration of 50ng/μl and incubation continued for 1 hour. The membrane was washed three times for 10 minutes each with binding buffer. The membrane was then incubated with binding buffer containing rabbit anti-BASP1 antibody for 30 minutes then washed three times for 5 minutes each. The membrane was then incubated with binding buffer containing anti-rabbit HRP for 30 minutes then washed three times for 5 minutes each and subjected to chemiluminescence. Cholesterol content of nuclear extracts was measured using an Amplex Red cholesterol assay kit (Invitrogen).

### Calcium Imaging

K562 cell line derivatives were washed with Tyrode’s solution (140 mM NaCl, 5 mM KCl, 1 mM MgCl_2_, 3 mM CaCl2, 10 mM Hepes, 10 mM glucose and 1 mM pyruvic acid (pH 7.4)). After loading with 2μM fura2/AM with Pluronic F-127 for 40 minutes, the cells were placed on cover slips and imaged on an Olympus IX71 microscope. Excitation wavelengths of 340 nm and 380nm were used with an emission wavelength of 510nm. Images were collected every 4s using Imaging Workbench 5.2 (Indec Biosystems). Cells were perfused with Tyrode’s solution and stimulated with 10 mM ATP. Imaging data were collected as a ratio of fluorescence intensities. An evoked response was defined as an increase in fluorescence that was greater than 2 S.D. values above baseline.

### Quantification and statistical analysis

Data are expressed as mean with standard deviation (SDM). Data distribution and significance between different groups was analyzed in Excel or OriginPro 7.5 using Student’s t test. Calcium imaging data were graphed without any filtering using OriginPro 7.5 software and statistical comparisons were made using a Fisher’s exact test.

## Acknowledgements

We are grateful to Adrian Bunzel for help with Fluorimetry. This work was funded by the BBSRC so SGER (BB/T001925/1), MRC to SGER (MR/K001027/1), NIH to KFM and SGER (1R01GM098609). AEL was supported by a Wellcome Trust PhD Studentship for the Dynamic Cell Biology program (083474).

## Author contributions

The study was conceptualized and designed by AEL, SC and SGER. All authors performed experiments and analysed the data. AEL and SGER wrote the manuscript. All authors discussed results from the experiments and commented on the manuscript. SGER supervised the project.

---

**Supplementary Figure 1.**
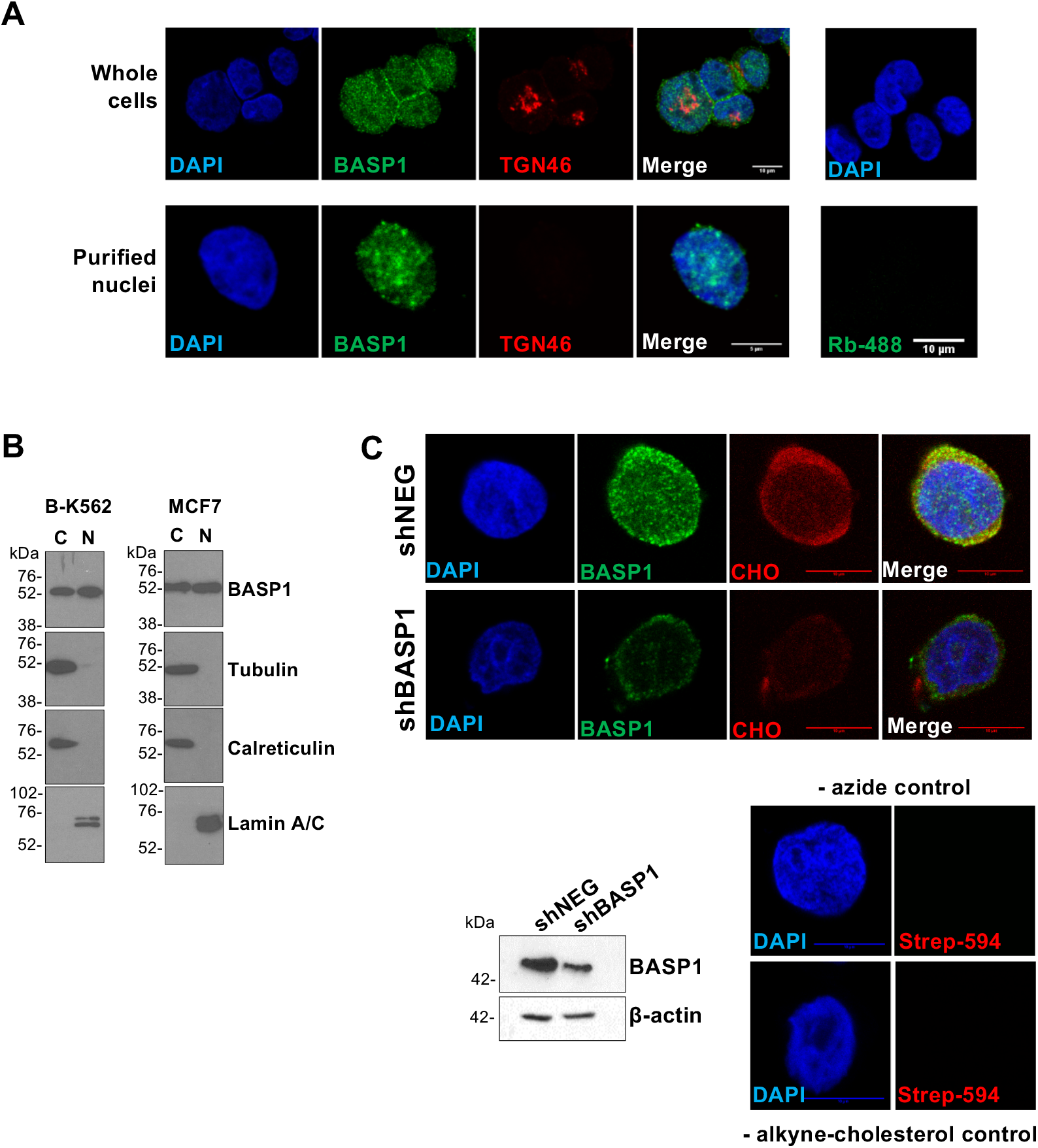
Analysis of Nuclear BASP1 and cholesterol. (A) Confocal images of whole K562 cells expressing BASP1 and purified nuclei immunostained for BASP1 and the golgi marker TGN46, plus DAPI. Scale bar 10μm in whole cells and 5μm in purified nuclei. (B) Cytoplasmic and nuclear extract were prepared from K562 and MCF7 cells and probed with antibodies to detect the proteins indicated at right. (C) MCF7 cells stably expressing a control shRNA (shNEG) or shRNA targeting BASP1 (shBASP1) were used to prepare nuclear extracts and immunoblotted with BASP1 or β-actin antibodies. Below, shNEG and shBASP1 MCF7 cells were incubated in lipid-free media with 10μg/ml alkyne-cholesterol. Nuclei were prepared followed by click chemistry and immunofluorescence with BASP1 antibodies and streptavidin antibodies to detect nuclear cholesterol (CHO).Scale bars are 10μm. At right are control immunofluorescence performed with nuclei purified from cells that were not incubated with alkyne-cholesterol.

**Supplementary Figure 2.**
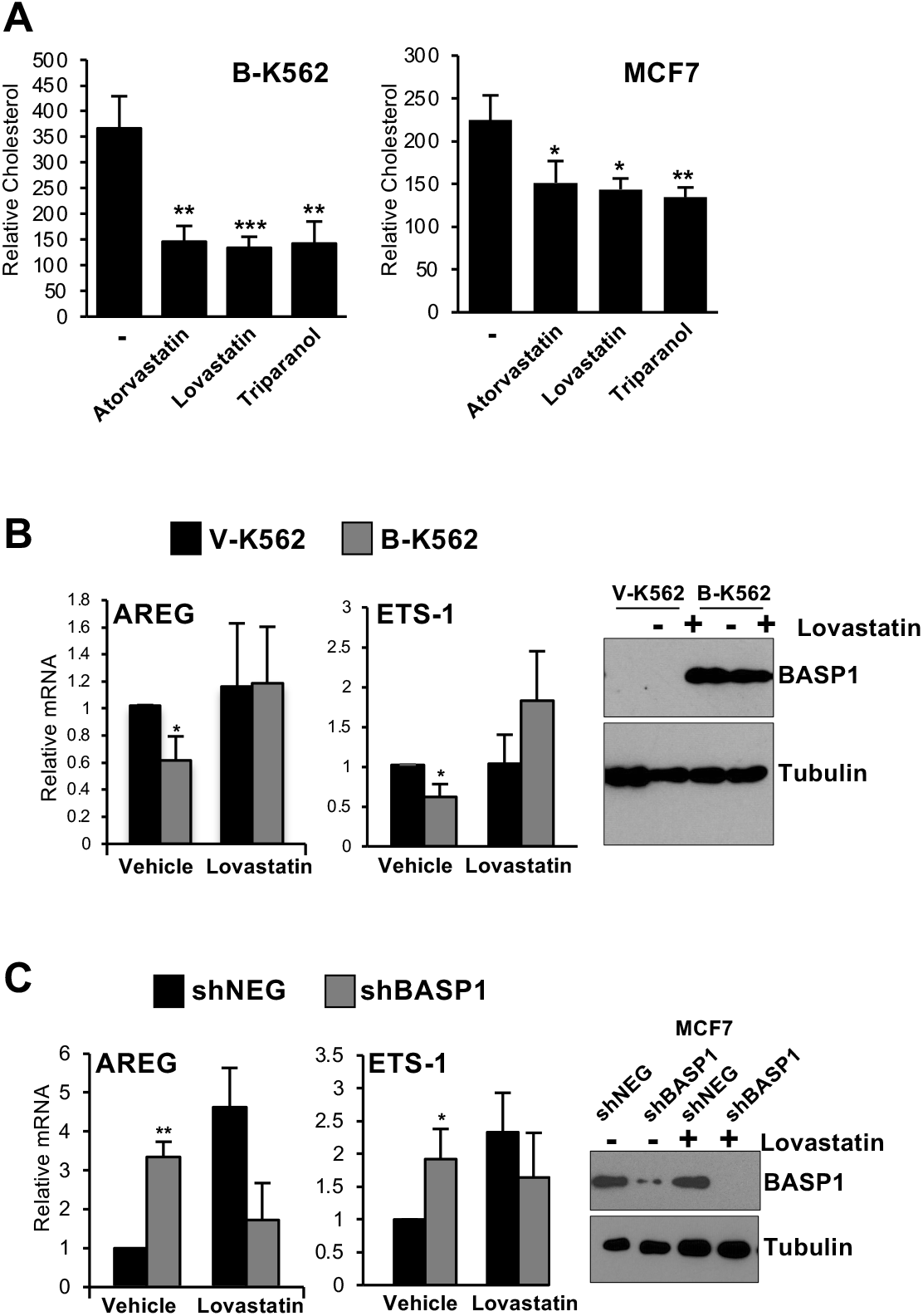
The effects of Statins on BASP1 transcriptional repressor function. (A) 10^6^ B-K562 or MCF7 cells were treated with either 20μM Atorvastatin, 50μM Lovastatin, 20μM Triparanol or drug vehicle (DMSO) for 48 hours, nuclear extracts prepared and assayed for cholesterol content. Results are expressed as relative cholesterol content/10^6^ nuclei. (B) V-K562 or B-K562 cells were treated with 50μM Lovastatin or drug vehicle (DMSO) for 48 hours. cDNA was prepared and expression of AREG and ETS-1 was determined relative to GAPDH. At right, a western blot of BASP1 and tubulin is shown. (C) MCF7 cells stably expressing a control shRNA (shNEG) or shRNA targeting BASP1 (shBASP1) were treated with 50μM Lovastatin or drug vehicle (DMSO) for 48 hours. cDNA was prepared and expression of AREG and ETS-1 was determined relative to GAPDH. At right, a western blot of BASP1 and tubulin is shown. Error bars denote SDM of at least 3 independent experiments (*p<0.05, **p<0.01, ***p<0.001 by student t-test).

**Supplementary Table 1.** Antibodies.

| **Antibody** | **Source** | **Cat. No.** | **RRID** |
| --- | --- | --- | --- |
| Normal rabbit IgG | Merck-Millipore | 12-379 | AB_145841 |
| Normal mouse IgG | Merck-Millipore | 12-371 | AB_145840 |
| Rabbit polyclonal anti H3K4me<sup>3</sup> | Merck-Millipore | 07-473 | AB_1977252 |
| Rabbit polyclonal anti H3K9Ac | Abcam | ab10812 | AB_297491 |
| Mouse monoclonal anti-RNA polymerase II | Abcam | ab 817 | AB_306327 |
| Mouse monoclonal anti-β-actin | Sigma Aldrich | A5316 | AB_476743 |
| Rabbit polyclonal anti-BASP1 | Carpenter et al., 2004 | doi: 10.1128/mcb.24.2.537-549.2004 |  |
| Sheep polyclonal anti-BASP1 | Carpenter et al., 2004 | doi: 10.1128/mcb.24.2.537-549.2004 |  |
| Mouse monoclonal anti-PIP2 | Abcam | ab11039 | AB_442848 |
| Rabbit polyclonal anti β-tubulin | Abcam | ab6046 | AB_2210370 |
| Rabbit polyclonal anti-calreticulin | Cell Signaling | 2891S | AB_2275208 |
| Mouse monoclonal anti-HA | Cell Signaling | 2367 | AB_10691311 |
| Anti-Mouse-HRP | Merck-Millipore | 71045-3 | AB_10808067 |
| Anti-Rabbit-HRP | Merck-Millipore | AP304P | AB_11212320 |

**Supplementary Table 2.**
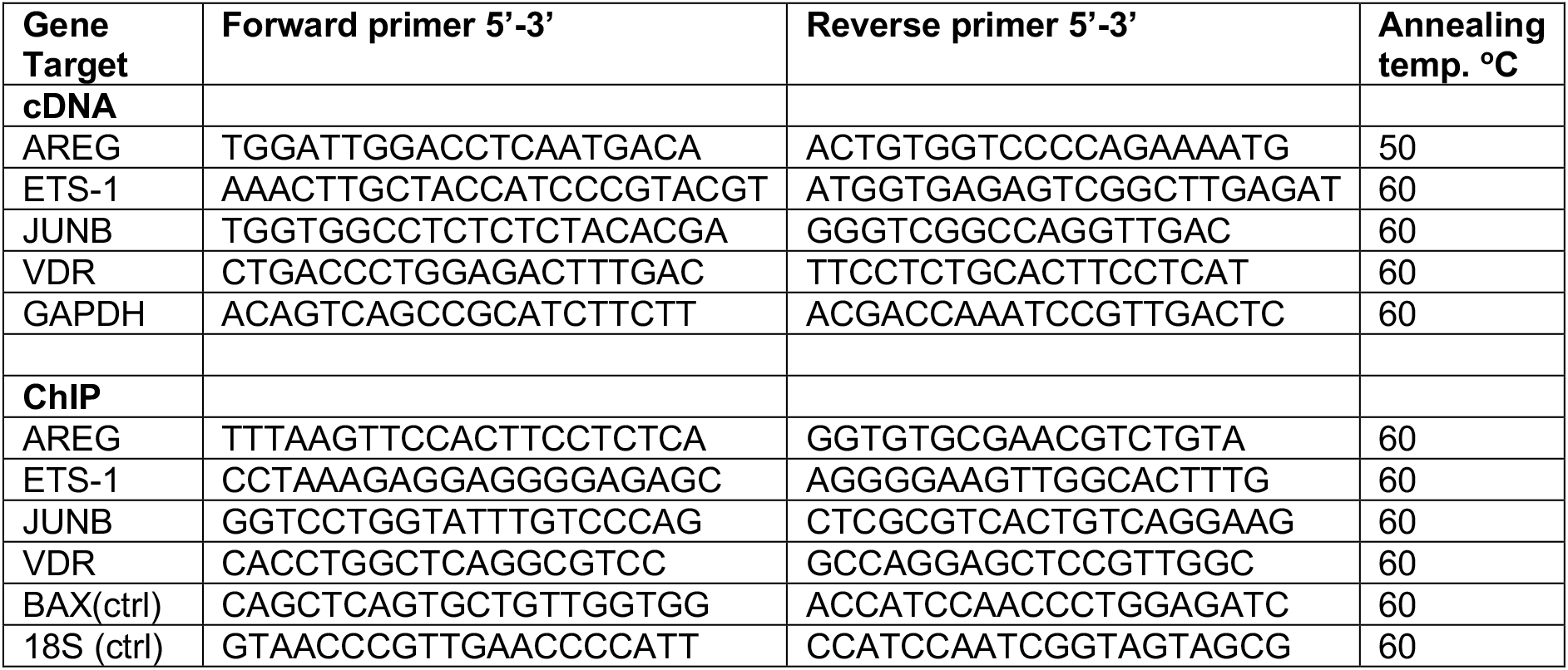
Primers for qPCR.

